# Evaluating bacterial and functional diversity of human gut microbiota by complementary metagenomics and metatranscriptomics

**DOI:** 10.1101/363200

**Authors:** Ravi Ranjan, Asha Rani, Patricia W. Finn, David L. Perkins

## Abstract

It is well accepted that dysbiosis of microbiota is associated with disease; however, the biological mechanisms that promote susceptibility or resilience to disease remain elusive. One of the major limitations of previous microbiome studies has been the lack of complementary metatranscriptomic (functional) data to complement the interpretation of metagenomics (bacterial abundance). The purpose of the study was twofold, first to evaluate the bacterial diversity and differential gene expression of gut microbiota using complementary shotgun metagenomics (MG) and metatranscriptomics (MT) from same fecal sample. Second, to compare sequence data using different Illumina platforms and with different sequencing parameters as new sequencers are introduced and determine if the data are comparable on different platforms. In this study, we perform ultra-deep metatranscriptomic shotgun sequencing for a sample that we previously analyzed with metagenomics shotgun sequencing. We validated the sequencing and analysis methods using different Illumina platform, and with different sequencing and analysis parameters. Our results suggest that use of different Illumina platform did not lead to detectable bias in the sequencing data. The analysis of the sample using MG and MT approach shows that some species genes are more highly represented in the MT than in the MG, indicating that some species are highly metabolically active. Our analysis also shows that ~52% of the genes in the metagenome are in the metatranscriptome, and therefore are robustly expressed. The functions of the low and rare abundance bacterial species remain poorly understood. Our observations indicate that among the low abundant species analyzed in this study some were found to be more metabolically active compared to others and can contribute distinct profiles of biological functions that may modulate the host-microbiota and bacteria-bacteria interactions.

## INTRODUCTION

The human microbiota represents a complex community of numerous and diverse microbes that is linked with our development, metabolism, physiology, health, and is considered functionally comparable to an organ of the human body (Cho & Blaser 2012, Human Microbiome Project 2012). Previous studies have established that a healthy human microbiota is associated with maintaining health, whereas dysbiosis has been associated with various pathologies and diseases such as obesity, inflammatory bowel disease, pulmonary diseases, urinary tract infection etc., (Iebba et al 2016, Pflughoeft & Versalovic 2012). Traditionally, identifying microbes relied on culture based techniques, however the majority (>90 – 95 %) of microbial species cannot be readily cultured using current laboratory techniques (Sharma et al 2005). Advancements in culture- and cloning–independent molecular methods, coupled with high-throughput next-generation DNA sequencing technologies have rapidly advanced our understanding of the microbiota. Additionally, with the rate of recent technological advancements, the DNA sequencing ventures have been introducing new DNA sequencers with versatile sequencing parameters. This has also complicated the comparison of data within and among the samples. Thus, there is a need to compare the sequencing data from same samples using different platforms. Many previous studies employed targeted amplicon sequencing of the conserved prokaryotic 16S ribosomal RNA (16S rRNA) gene (Human Microbiome Project 2012, Huse et al 2012, Stulberg et al 2016). This method identifies operational taxonomic units (OTUs) and are correlated with bacterial taxa; however, assignment of taxa defined by OTUs is commonly limited to the genus level due to low accuracy at the species level. In contrast, metagenomics shotgun sequencing (MGS), which is employed in our study, can determine taxonomic annotations at the species level.

Although the association of multiple diseases with dysbiosis of the microbiome has been established, the elucidation of the underlying biologic mechanisms that promote pathological phenotypes has been elusive in most cases. A major limitation of both targeted amplicon and metagenome shotgun sequencing is that bacterial functions are predicted based on the genome sequence of the associated taxa. However, it is well established that there is differential bacterial gene expression at the transcriptional level in response to environmental and dietary exposures. For example, it has been reported that there is a set of constitutively expressed core genes that mediate core microbial functions as well as a highly regulated subset of genes that respond to unique environmental influences (Booijink et al 2010, Ursell & Knight 2013). In addition, some bacteria may exist in an inert state or spore form and thus not contribute to the biological response (Franzosa et al 2014). Thus, an analysis of bacterial gene expression with metatranscriptomics approach could provide additional insight into the biological functions of specific microbiomes.

The gut microbiota is composed of highly abundant few species and less abundant many rare bacterial species, thus to understand the complex functions of the microbiota it is essential to understand the functions of both the high- and low-abundant bacterial species. Analyses of MG and MT data are often challenged by the sequencing depth, parameters, and sequencing platforms, which limits the power of functional classification and abundance estimation, this in turn hampers the downstream data analyses of differentially expressed genes. The unique feature of our study is that we are comparing the sequencing reads at different depths, platform, read length, read and contig based comparison for MG and MT for the same sample. To develop a comprehensive understanding of the ecological functions of a microbiome, it is essential to determine not only the metatranscriptome, but also to ascertain the functional contributions of both the abundant and the rare species in a microbiome. To investigate these questions, we analyzed both the metagenome and the metatranscriptome using shotgun sequencing which can determine the abundance of gene transcripts relative to the abundance of the genome. This allowed us to identify both over- and under-expressed transcripts. In this study, we identified biological functions in both rare and abundant bacterial species using metagenomic and metatranscriptomic methods optimized and validated in our laboratory.

## MATERIAL AND METHODS

### Subject recruitment and sample collection

The study was approved by the Institutional Review Board of the University of Illinois at Chicago, and the experimental methods were performed in accordance with the approved guidelines. A 33 year, male subject without known medical conditions provided the signed informed consent and self-collected stool in a EasySampler Stool Collection kit (Alpco Diagnostics). The fecal sample was immediately aliquoted into sterile 1.5 ml Eppendorf safe-lock tubes and stored at −80°C till further DNA and RNA isolation was carried out.

### RNA isolation from fecal sample and mRNA enrichment

The objective of the study was to perform matched metagenome and metatranscriptome studies of the same fecal sample. We investigated the same fecal sample we previously analyzed by metagenomics sequencing. Total RNA was isolated using the PowerMicrobiome RNA Isolation Kit (Catalog # 26000-50, MO BIO Laboratories, Inc) from a fecal sample. For efficient lysis of the microbes in the sample, 200 μL of Phenol/Chloroform/Isoamyl alcohol (25:24:1) (Catalogue #327115000, Acros Organics) was added to the reagents provided with the kit. The contents were vortexed for 1-2 min with a table top vortexer and homogenized twice at speed 10 for 5 min with air-cooling using the Bullet Blender Storm Homogenizer (Catalogue # BBY24M, Next Advance Inc). Total RNA was isolated with the manufacturer’s recommended procedure including the on-column DNase treatment (to remove the potentially co-isolated DNA). The RNA was eluted with 1×TE, pH 8.0, and stored at −80°C. The quality and quantity of the DNA was accessed using a spectrophotometer (NanoPhotometer Pearl, Denville Scientific, Inc), agarose gel electrophoresis, fluorometer (Qubit^®^ RNA Broad Range assay, Life Technologies Corporation), and Agilent RNA 6000 Nano Kit on 2100 Bioanalyzer instrument (Agilent Technologies, Inc.). Total RNA was enriched for mRNA by subtractive hybridization using the MICROBExpress^™^ Bacterial mRNA Enrichment Kit following manufacturers recommended protocol (Ambion, Life Technologies). The mRNA enrichment and rRNA depletion was analyzed using an Agilent RNA 6000 Nano Kit on 2100 Bioanalyzer instrument (Agilent Technologies, Inc.).

### Fecal metatranscriptome library preparation and shotgun sequencing

The enriched mRNA was mechanically fragmented to a size range of ~200 bp with an ultrasonicator using the adaptive focused acoustics with the following manufacturer recommended protocols (Covaris S220 instrument, Covaris Inc). The fragmentation of mRNA was assessed using Agilent RNA 6000 Pico Kit on 2100 Bioanalyzer instrument (Agilent Technologies, Inc). The metatranscriptome libraries were prepared using NEBNext Ultra RNA Library Prep Kit for Illumina (New England BioLabs Inc). The quality and quantity of all the final libraries were analyzed with an Agilent DNA 1000 Kit on the 2100 Bioanalyzer Instrument and Qubit. The final libraries were quantitated and validated by qPCR assay using the PerfeCTa NGS Library Quantification Kit for Illumina (Quanta Biosciences, Inc.) using the CFX Connect Real-Time PCR Detection System (Bio-Rad Laboratories, Inc). Sequencing of one of the MT library was performed on a Illumina HiSeq 2000 using the TruSeq SBS v3 reagent for paired-end 100 read length (BGI Americas) (labeled as HS100), and on Illumina MiSeq using v3-600 cycle kit for paired-end 301 bases (labeled as MS301). Another set of twelve libraries was sequenced on Illumina MiSeq using 151 paired end chemistry (labeled as MS151). Manufacturer’s recommended protocol was used for performing the sequencing reaction on both the HiSeq and MiSeq platforms.

### Data analysis

The individual twelve libraries were analyzed for taxonomic and functional annotation, also all of the 12 sequence files were combined in silico and were labeled as (MS151)-Lib-All. The sequence files (HS100, 12(x) MS151, and MS301) were combined in silico and labelled as HS100+MS151+MS301.: The sequence reads were processed and analyzed using the CLC Genomics workbench version 7.5 (Qiagen, Aarhus, Denmark). Raw reads were trimmed to a minimum Phred quality score of 20. Raw reads were filtered by mapping against human reference genome to remove human sequences. The non-human reads were de novo assembled using the CLC assembler using a word size (k-mer) of 50, minimum contig length 200bp, to construct the de bruijn graphs. De novo assembly was used to map reads back to the contigs (mismatch cost 2, insertion cost 3, deletion cost 2, length fraction 0.8, similarity fraction 0.8). Taxonomic and functional annotations of the reads and contigs were obtained using the automated annotation pipeline at MG-RAST web server using the default parameters (best hit classification, maximum e-value 1e-5 cutoff, and minimum 60% identity cutoff) using M5NR and KEGG databases (Meyer et al 2008, Mitra et al 2011). The limma analysis was used to identify species and KEGG functional pathways that were differentially abundant between metagenome (MG) and metatranscriptome (MT) (Praveen et al 2015). Limma uses an empirical Bayes method to test the differential expression based on the fitting of each species/gene to a linear model (Smyth 2004). This provides the rich features for complex experimental designs and overcomes the small sample size problem, in addition to providing enhanced biological interpretation for co-regulated sets of genes (Ritchie et al 2015). A *p* value cutoff of 0.05 after multiple testing correction based on Benjamini-Hochberg method (Benjamini & Hochberg 1995), and a log_2_ fold change ≥1 were used to select the differentially abundant species and pathways. The data files were visualized in MeV v 4.9.0 (TM4, Boston, MA, USA) (Saeed et al 2003). The metatranscriptome data was used to compare with the previously reported metagenome data of the same sample from our group (Ranjan et al 2016).

## RESULTS

### Ultra-deep metatranscriptomic shotgun sequencing (MTS)

In our previous study of ultra-deep metagenome shotgun sequencing (MGS) we demonstrated effective identification of abundant species (defined as >1% relative abundance) with as few as 500 reads; however, the detection of low abundance or rare species required high numbers of sequence reads. For example, with a total of 163.7 million sequence reads generated by metagenome shotgun sequencing (MGS), the rarefaction curve did not show saturation for the identification of additional species (Ranjan et al 2016). Based on these data, in the current study of the metatranscriptome we performed ultra-deep MTS sequencing. We performed optimization and validation of our sequencing protocol using multiple sequencing platforms and analytic strategies (Fig. 1). High quality total RNA was isolated (Supplementary Fig. 1A), and the bacterial mRNA was enriched from the total RNA using subtractive hybridization, which depleted most of the rRNA (Supplementary Fig. 1B). The enriched mRNA was mechanically fragmented and libraries were constructed (Supplementary Figs. 1C and1D). To evaluate technical reproducibility, we constructed 12 unique indexed metatranscriptome libraries from a single fecal sample. High quality libraries were prepared for sequencing on Illumina’s MiSeq and HiSeq 2000 platforms (Supplementary Fig. 1E). We obtained from 3.6 to 5.4 million high quality sequence reads for the 12 replicate libraries sequenced on MiSeq for 151 PE and 32.7 to 56.5 million reads on a HiSeq 2000 platform using 100 and 151 PE sequencing parameters. In total, we obtained a total of 139.6 million sequence reads by combining the HiSeq and MiSeq sequence data in silico (HS100+MS151+MS301) (Table 1).

**Figure 1.**
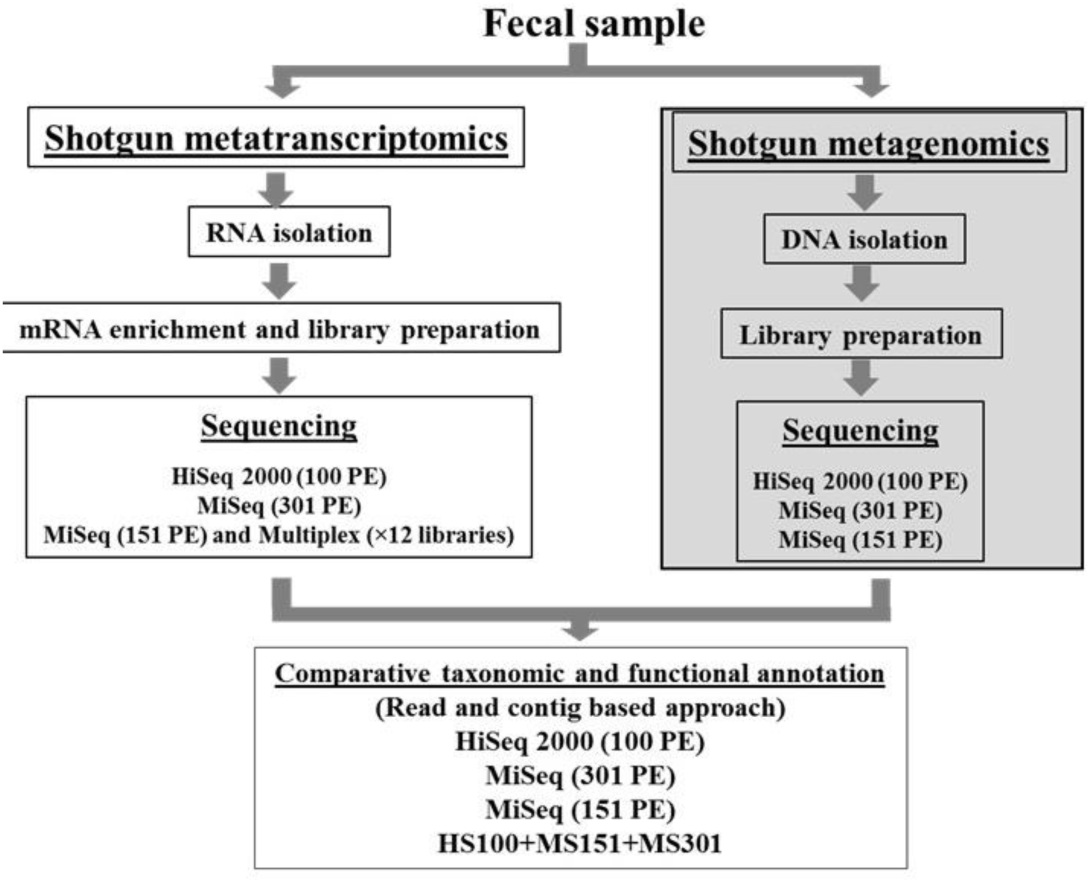
Experimental strategy to compare the metatranscriptome and metagenome using multiple Illumina sequencing platforms and data analysis. Schematic for metagenome and metatranscriptome sequence analysis by shotgun sequencing approach. The shotgun sequencing was performed using Illumina HiSeq 2000 (100 paired-end), and Illumina MiSeq (151 and 301 paired-end). The data was analyzed by read and contig based approach using the MG-RAST. Note that the metagenome data has been published (Ranjan et al., 2016), represented in shaded box.

**Table 1.** Metatranscriptome sequence statistics.

| Sample name | Read |  |  | Contig |  |  |
| --- | --- | --- | --- | --- | --- | --- |
|  | Number of PE reads (M) | Average length (bp) | Total bases (Mb) | % of reads assembled in contig | Number of contig | Average length (bp) |
| MS151_library 1 | 4.8 | 145 | 690.5 | 98.4 | 7,253 | 207 |
| MS151_library 2 | 4.8 | 143 | 690.2 | 98.2 | 7,517 | 209 |
| MS151_library 3 | 5.4 | 143 | 765.2 | 98.4 | 8,291 | 202 |
| MS151_library 4 | 4.5 | 143 | 649.9 | 98.4 | 7,364 | 209 |
| MS151_library 5 | 4.8 | 144 | 686.8 | 98.0 | 7,072 | 212 |
| MS151_library 6 | 4.3 | 145 | 625.0 | 98.0 | 6,183 | 213 |
| MS151_library 7 | 4.9 | 143 | 696.9 | 98.4 | 7,635 | 204 |
| MS151_library 8 | 3.6 | 147 | 525.9 | 97.3 | 5,889 | 222 |
| MS151_library 9 | 4.9 | 146 | 707.4 | 97.9 | 7,875 | 208 |
| MS151_library 10 | 4.5 | 145 | 653.9 | 98.4 | 6,779 | 215 |
| MS151_library 11 | 4.8 | 145 | 698.7 | 98.1 | 7,117 | 211 |
| MS151_library 12 | 5.3 | 145 | 768.9 | 98.0 | 8,249 | 212 |
| HS100 | 50.4 | 100 | 5,039.2 | 98.6 | 11,713 | 203 |
| MS151 (Lib-all) | 56.5 | 144 | 8,159.3 | 99.3 | 56,491 | 208 |
| MS301 | 32.7 | 178 | 5,837.7 | 98.3 | 108,905 | 314 |
| HS100+MS151+MS301 | 139.6 | 136 | 19,036.1 | 99.1 | 216,712 | 268 |

HS100: HiSeq 2000 - 100 PE; MS151: MiSeq - 151 PE; MS151: MiSeq - 301 PE; M: Million; bp: basepair; Mb: Mega bases; PE: Paired-end sequencing.

### Comparison of analytic strategies

In our previous analysis of ultra-deep MGS data, we observed a substantial increase in the average length of the assembled contigs (904 bp) compared with the average read length 170 bp., and the average N50 length of the contigs was 6,262 bp (Ranjan et al 2016). Therefore, we compared the effect of analyzing the reads versus assembled contigs in the metatranscriptome (MT) data. In the MT data, the average contig length was 268 bp which was modestly longer than the average read length of 136 bp (Table 1). The short length of the assembled MT contigs compared to the metagenomic (MG) contigs is likely due to the smaller size of the microbial transcripts compared to the larger size of the genomes. In terms of reproducibility, we did not detect significant differences between the number of reads or assembled contigs among the 12 replicate libraries as analyzed by *Shapiro-Wilk* normality test (data not shown). Thus, the assembly of the contigs generated a modest increase in length compared with average read length of the MT reads.

Next, we compared the bacterial taxonomic assignments based on read and contig analyses. Analysis at the phyla, genera and species levels all demonstrated the reproducibility of the replicate libraries, respectively (Supplementary Figs. 2-4). However, we detected differences in the relative abundance of specific taxa in the read and contig based analyses. Thus, the taxonomic identification was inconsistent between read and contig based analysis at both phylum and genus level. For example, we observed an increase in the Bacteroidetes and decrease in Firmicutes with the contig analysis. Differences in relative abundance in the MT data were also observed at the genus and species levels. There were 21 and 11 genera, and 22 and 19 species in the read and contig based analysis that were above 1% abundance, respectively (Supplementary Figs. 3 and 4, Supplementary Tables 1 and 2). We further analyzed the bacterial diversity of combined MT datasets (HS100, MS151, MS301 and HS100-MS151-MS301) to increase the sequencing depth and coverage. We find similar observations in the distribution of bacterial phyla (Supplementary Fig. 5A). We observe that the increase in number of reads resulted in increase of depth of coverage, whereas no significant increase in contig length was detected. In summary, we previously showed that a contig based analysis is more specific for species identification (Ranjan et al 2016) in the MGS dataset; however, these data suggest that a read based analysis is more comprehensive for identification of both genera and species in metatranscriptome data.

To determine if different numbers of reads were skewing the analyses, we generated datasets that contained an equal number of reads. We randomly sampled 30 million reads from the HiSeq 100 PE, MiSeq 151 PE (MS151) and MiSeq 301 PE (MS301) data, and the reads were assembled into contigs. More contigs were generated in MS301 (97,631) compared to HS100 (8,253) and MS151 (42,153), most likely because of a longer sequencing read length. However, there was no substantial increase in the average length of contigs most likely due to the limitation based on transcript length (Supplementary Table 3). We observed a similar abundance profile of bacterial phyla, genera and species as in the complete datasets indicating that differences in read number were not skewing the assignment of taxa in the contig analyses (Supplementary Fig. 5B, and Supplementary Table 4).

### Comparison of the metatranscriptome with the metagenome

In total, we identified 1,888 and 1291 bacterial species in the metagenome (MG) [MG-HS100-MS151-MS301, (Ranjan et al 2016)], and the metatranscriptome (MT) (MT-HS100-MS151-MS301) data, respectively (Fig. 2A). 1245 bacterial species were shared among the MG and MT (Fig. 2A), representing the metabolically active species, in the sample at this particular time point. In the phylum Firmicutes, Bacteroidetes, Actinobacteria, Proteobacteria, Fusobacteria, and Verrucomicrobia 356, 117, 138, 439, 23, and 6 species were shared, respectively. This accounted for 60% to 92% of the species shared between the MG and MT defined phyla (Fig. 2B). The detection of MG sequences lacking corresponding MT reads suggests unexpressed genes or even dormant bacteria. As expected, very few sequences were unique to the MT, and they were present in extremely low abundance (< 0.001%) presumably because transcripts are not expressed in the absence of the genome, and likely these sequences were not identified in MG because of relatively low abundance (Supplementary Table 6). Most (50%) of the sequences identified in the phylum proteobacteria were closely related to uncultured bacterial sequences. To determine the relative transcriptional activity of individual taxa and individual genes, we compared the relative abundance in the combined MT data (HS100-MS151-MS301) to our previously reported MG data for the same sample (Ranjan et al 2016). In an analysis of the MT at the phyla level, we observed that the abundance of Bacteroidetes transcripts was high, whereas the abundance of transcripts representing Firmicutes, Actinobacteria, Fusobacteria, and Verrucomicrobia was low. This was observed across all the sequencing platforms and read lengths (Fig. 2C). The abundance of the Fusobacteria and Verrucomicrobia was approximately 100-fold lower than the other Phyla (note Y-axis scale).

**Figure 2.**
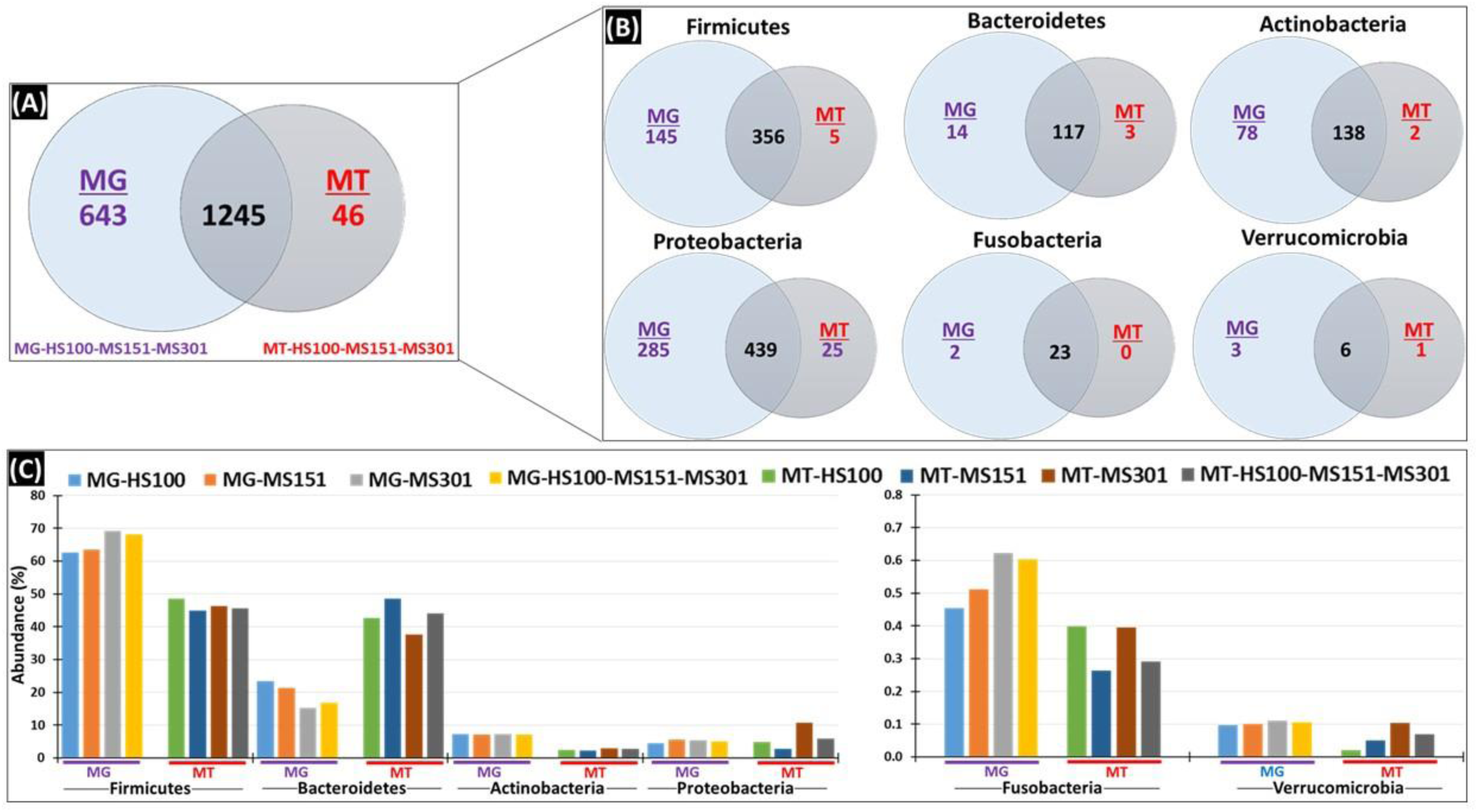
Taxonomic analysis: Comparison of metagenome (MG) and Metatranscriptome (MT). The MG and MT sequence obtained after sequencing using platforms (HS100, MS151 and MS301) were assembled into contig and were analyzed for taxonomic annotation. (A) The total bacterial species in MG-HS100-MS151-MS301 and MT-HS100-MS151-MS301 data. (B) Bacterial species in MG-HS100-MS151-MS301 and MT-HS100-MS151-MS301 in different phyla. (C) The abundance of bacterial phyla in MG and MT with different sequencing parameters - Firmicutes, Bacteroidetes, Actinobacteria, Proteobacteria, Fusobacteria and Verrucomicrobia.

### Analysis of predicted biological functions

We analyzed the functional profiles based on gene expression in the metatranscriptome using the MG-RAST KEGG annotation suite. KEGG annotates functions from level 1 through 4 with level 1 containing the most general categories and level 4 the most specific (Mitra et al 2011). We analyzed the data for biological functions at all four levels. Of note, a similar relative abundance of the functions was detected at levels 1 to 4 among the both read and contig based analysis, respectively (Supplementary Figs. 6-9), although minor differences were detected in the abundances of some functions at levels 3 and 4. We observe the similar distribution trends in the 30 million randomized MT reads and the assembled contigs (Supplementary Fig. 10). This implies that the identified functions are similar in either the read or contig based analysis of the MT data with slight variations.

We investigated the MG and MT data at the species level. Interestingly, we observed that few of the species (for example, *Faecalibacterium prausnitzii, Bacteroides spp., B. thetaiotaomicron, B. vulgatus, B. ovatus* among others) had a higher relative representation in MT than MG, indicating that these species are highly transcriptionally active (Fig. 3 and Supplementary Fig. 11). However, the species *B. fragilis* did not have increased transcriptional activity as compared to other *Bacteroides spp*. As shown in a scatter plot, *F. prausnitzii*, *Bacteroides spp*., and *Alistipes putredinis* were highly transcriptionally active at a significant level (log fold difference ≥3, *p* adj. <0.05) whereas *Clostridium saccharolyticum, Eubacterium rectale* and *Ruminococcus obeum* (log fold difference ≥-1, *p* adj. <0.05) were low in transcriptional activity (Fig. 4A).

**Figure 3.**
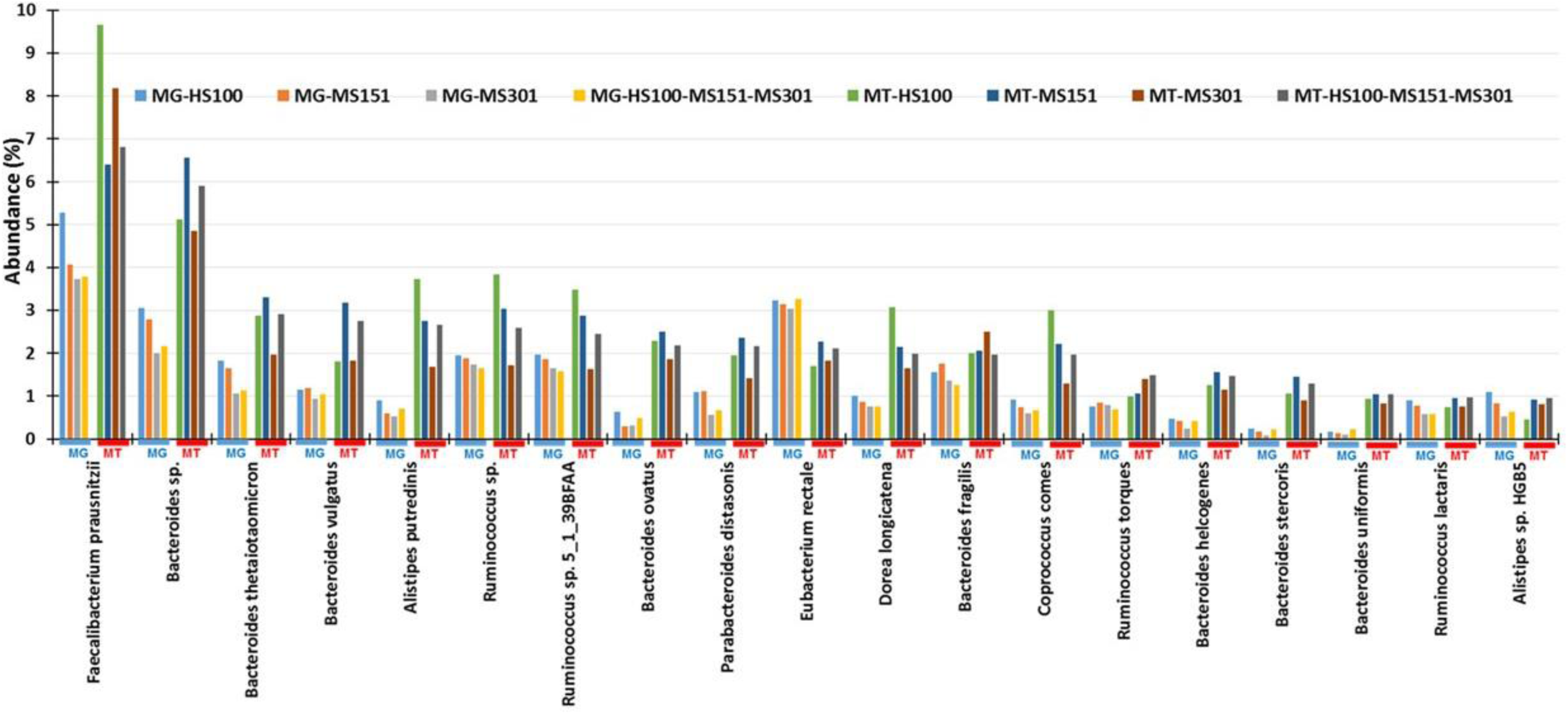
Abundance of bacterial species in metagenome and metatranscriptome. Bacterial species above 1% (sorted high to low) are shown in MT-HS100-MS151-MS301.

**Figure 4.**
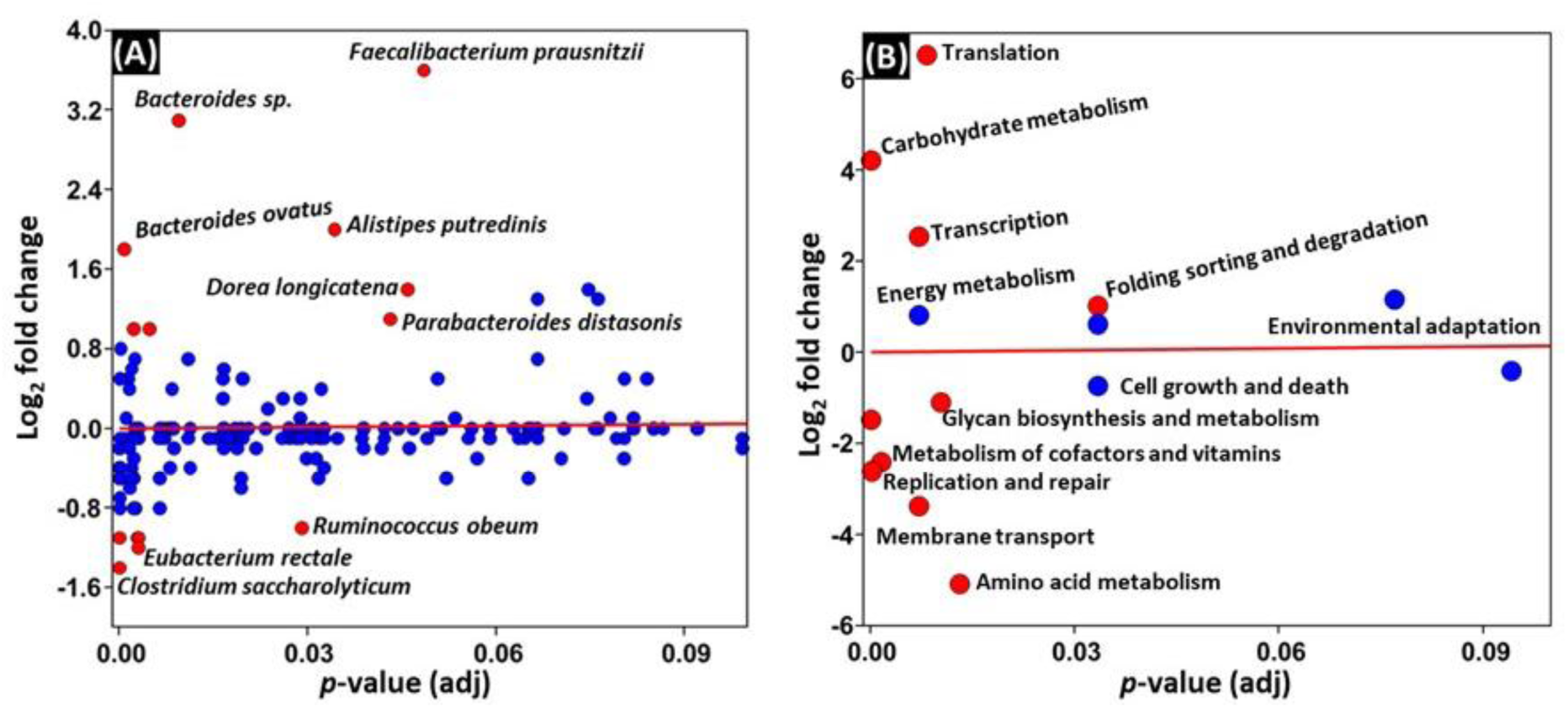
Differential abundant species and KEGG functional categories. The scatter plot for differential abundant bacterial species (A) and differentially predicted and expressed KEGG functional categories (B) in the metagenome and metatranscriptome. A *p* value cutoff of 0.05 (after FDR correction based on Benjamini-Hochberg method) and a log fold change ≥1 were used to select the differentially abundant species and functional categories. Significant values for different species and pathways are shown in red and non-significant values are shown with blue circles.

We compared the abundance of KEGG functions detected in the MT data to the predicted functions in the MG data. The analysis revealed that genes involved in translation, carbohydrate metabolism, and transcription were highly abundant in MT (log_2_ fold change >3, *p*< 0.05), compared to low abundance of glycan biosynthesis and metabolism, metabolism of cofactors and vitamins, replication and repair, membrane transport and amino acid metabolism (log_2_ fold change >-2, *p* adj. < 0.05) (Fig. 4B). Translation and amino acid metabolism showed the largest differential expression with a fold change of >±5 (*p* adj. <0.05), respectively. We observed similar patterns at the more specific levels 2, 3 and 4 (Supplementary Fig. 12-15). In this fecal sample, in total we detected 1916 functions at KEGG level 4 assignments in MG, compared to 1067 in MT. The MG and MT data shared 52% (1014) of the total functions, revealing the shared functional genes involved in active physiological functions of the gut microbiota which can be detected in MG and MT in a given time point (Fig. 5). Our analysis indicated that MG and MT overlapping genes are metabolically active genes. Genes which are only detected in the MT are even more metabolically active. On the other hand, if genes were detected only in MG and not in the MT, this may also suggest that genes may be present but not active in a given time.

**Figure 5.**
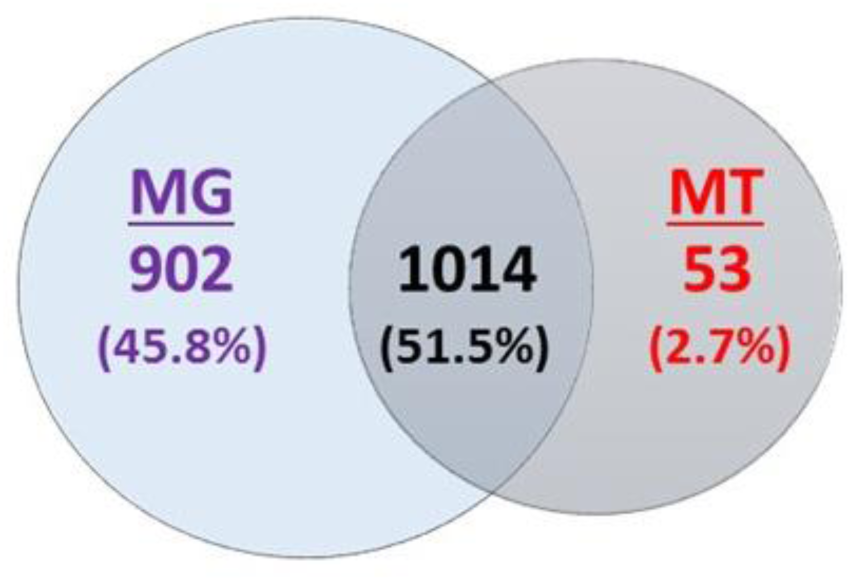
Comparison of the metabolic functional of metagenome (MG) and metatranscriptome (MT). Venn diagram for unique and shared metabolic functions identified by KEGG at functional level 4 in the MG (MG-HS100-MS151-MS301) and MT (MT-HS100-MS151-MS301).

### Contribution of functions in the metatranscriptome by individual bacterial phylum

We further explored the functional contribution of the gut microbiota at the individual phylum level comprising of Firmicutes, Bacteroidetes, Actinobacteria, Proteobacteria, Fusobacteria and Verrucomicrobia, as these are abundant in the gut. There were differences in the expression of the genes in each phylum (Supplementary Figs. 16-18). At the KEGG Level 1 functional category, 50% of the functions were related to metabolism in each phylum (Firmicutes, Bacteroidetes, Actinobacteria, Proteobacteria, Fusobacteria and Verrucomicrobia), followed by genetic and environmental information processing functional categories. Of note few functional categories related to the phylum Fusobacteria and Verrucomicrobia were detected (Supplementary Fig. 16). We further focused our analysis on Fusobacteria and Verrucomicrobia, as these phyla are present in low abundance (<1% and <0.1% abundance, respectively) and not well characterized in the gut microbiota (Fig. 2C).

In phyla - Firmicutes, Bacteroidetes, Actinobacteria, and Proteobacteria, the genes involved in carbohydrate metabolism were abundant, followed by amino acid metabolism and translation. There were no translation and/or transcription functions detected in Fusobacteria and Verrucomicrobia (Supplementary Fig. 17). However, Fusobacteria and Verrucomicrobia contributed towards the expression of specific genes involved in carbohydrate and amino acid metabolism pathways compared to other phyla (Figs. 8 and 9, Supplementary Fig. 18). For example, the genes *glgB* (1,4-alpha-glucan branching enzyme), *pgi* (glucose-6-phosphate isomerase) involved in starch, and sucrose metabolism and glycolysis/gluconeogenesis were highly expressed by Fusobacteria (Supplementary Fig. 18). Also, the genes involved in oxidative phosphorylation such as *atpD* (F-type H+-transporting ATPase subunit beta), *ppa* (inorganic pyrophosphatase) and *nuoE* (NADH-quinone oxidoreductase subunit E) were also enriched in Fusobacteria (Figs. 6, and Supplementary Fig. 18). On the other hand, the phylum Verrucomicrobia was enriched for genes invloved in alanine, aspartate and glutamate metabolism [*gdhA:* glutamate dehydrogenase (NADP+), *purB*: adenylosuccinate lyase], ABC transporters [*msmX*: maltose/maltodextrin transport system ATP-binding protein] and amino sugar and nucleotide sugar metabolism [*npdA*: NAD-dependent deacetylase] (Fig. 7 and Supplementary Fig. 18). These results show the high abundance of transcripts contributed by the rare abundant bacterial species in the community may contribute unique biological functions to the microbiome that have the potential to affect the host physiology.

**Figure 6.**
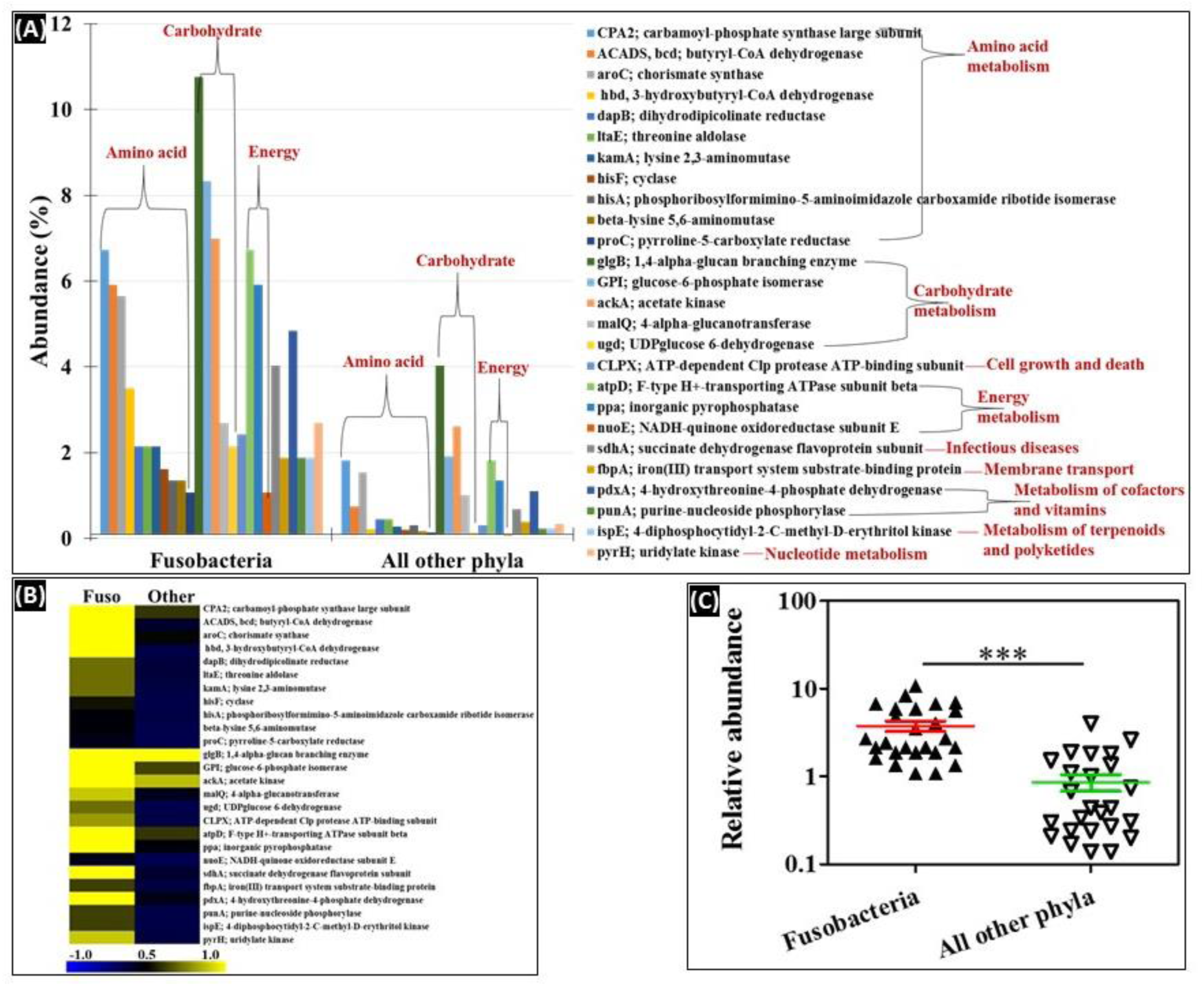
Metatranscriptome analysis of phylum Fusobacteria. (A) Relative abundance of Fusobacteria genes compared to all other phyla. (B) Heat-map representation of the genes. The color scheme represents the range of gene abundance values based on Spearman Rank correlation. C) Significant difference in log abundance of genes highly abundant in Fusobacteria compared to all other phyla. *p*<0.05, Mann-Whitney *U* test. Other phyla include Firmicutes, Bacteroidetes, Proteobacteria, Actinobacteria and Verrucomicrobia.

**Figure 7.**
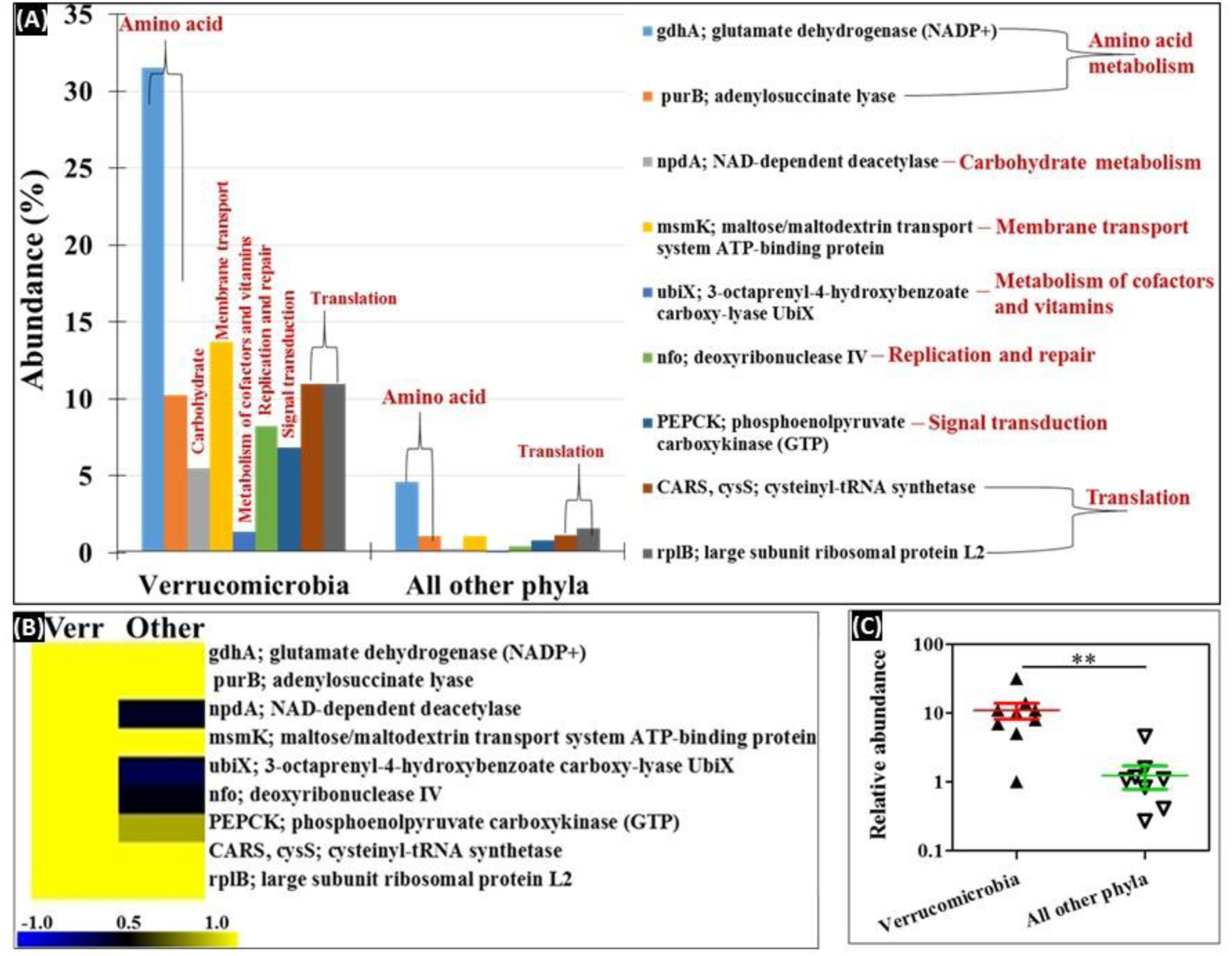
Metatranscriptome analysis of phylum Verrucomicrobia. (A) Relative abundance of Verrucomicrobia genes compared to all other phyla. (B) Heat-map representation of the genes. The color scheme represents the range of gene abundance values based on Spearman Rank correlation. (C) Significant difference in log abundance of genes highly abundant in Verrucomicrobia compared to all other phyla. *p*<0.05, Mann-Whitney *U* test. Other phyla include Firmicutes, Bacteroidetes, Proteobacteria, Actinobacteria and Fusobacteria.

### Diversity analysis of bacterial species and functions

The Shannon diversity index for estimating the bacterial diversity in MG (5.4 ± 0.1) and MT (4.9 ± 0.1) was significantly different (*p*<0.05), however no significant difference was observed in species evenness (0.7 ± 0.0). Similarly, the index for diversity of functional genes in MG (6.7 ± 0.0) and MT (6.0 ± 0.3) was significantly different (*p*<0.05), also a significant difference was observed in functional evenness in MG (0.89 ± 0.01) and MT (0.93 ± 0.01). The Shannon diversity index analysis at both taxonomic and functional level indicated that the MG was more diverse than the MT, most likely due to unexpressed genes or dormant bacteria (Supplementary Fig. 19).

### Mapping the genomic and transcriptomic KEGG pathways

We mapped the predicted (MG) and expressed (MT) functions onto pathways using KEGG Mapper suite. Almost all (more than 99%) of the functions identified by MT were also identified in MG (Fig. 8 and Supplementary Fig. 20). However, some functions were identified only in the MG dataset suggesting that not all of the predicted functions in the metagenome are expressed, which supports the notion that the metagenome may not be an accurate proxy of microbiota function. The genes are in the (meta)genomes; they could be expressed under different conditions; therefore, they define the functional potential of the organisms. Linear regression analysis was applied to the MT and MG data examined from the perspective of species and function. The linear regression analysis at the species level was correlated among the MG and MT and 58% of the variation in the MT can be explained by the species composition of the MG (Spearman’s *r* = 0.83; *r*^2^=0.58=58%) (Fig. 9A). A similar correlation was observed at functional level 4 in MG and MT (Spearman’s *r* = 0.76; *r*^2^=0.53=53%) (Fig. 9B). In other words, more than 50% of the variation in the microbial community MT can be explained by MG composition at species level, or conversely, approximately 50% of transcriptional activity is regulated and presumably dependent on host or environmental factors.

**Figure 8.**
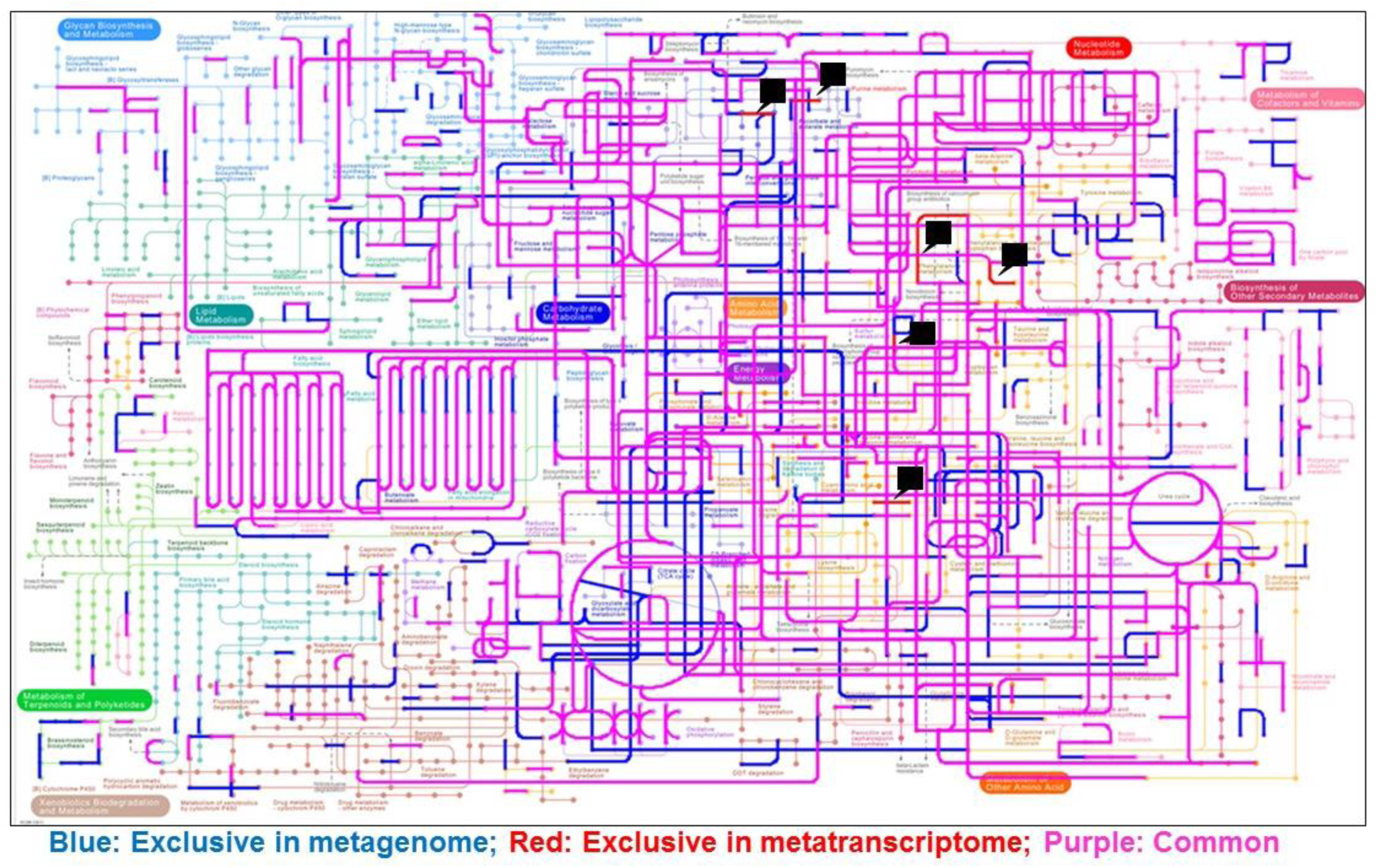
Differential metabolic gene expression. Metabolic pathway reconstruction in metagenome and metatranscriptome were analyzed using the KEGG mapper. Functions identified in the metagenome (MG-HS100+MS151+MS301) and metatranscriptome (MT-HS100+MS151+MS301). Blue: predicted functions exclusive in metagenome; Purple: Common in metagenome and metatranscriptome; Red: Exclusive in metatranscriptome. Black arrow head represents the functions in MT. Function in individual data are shown in Supplementary Fig. 20.

**Figure 9.**
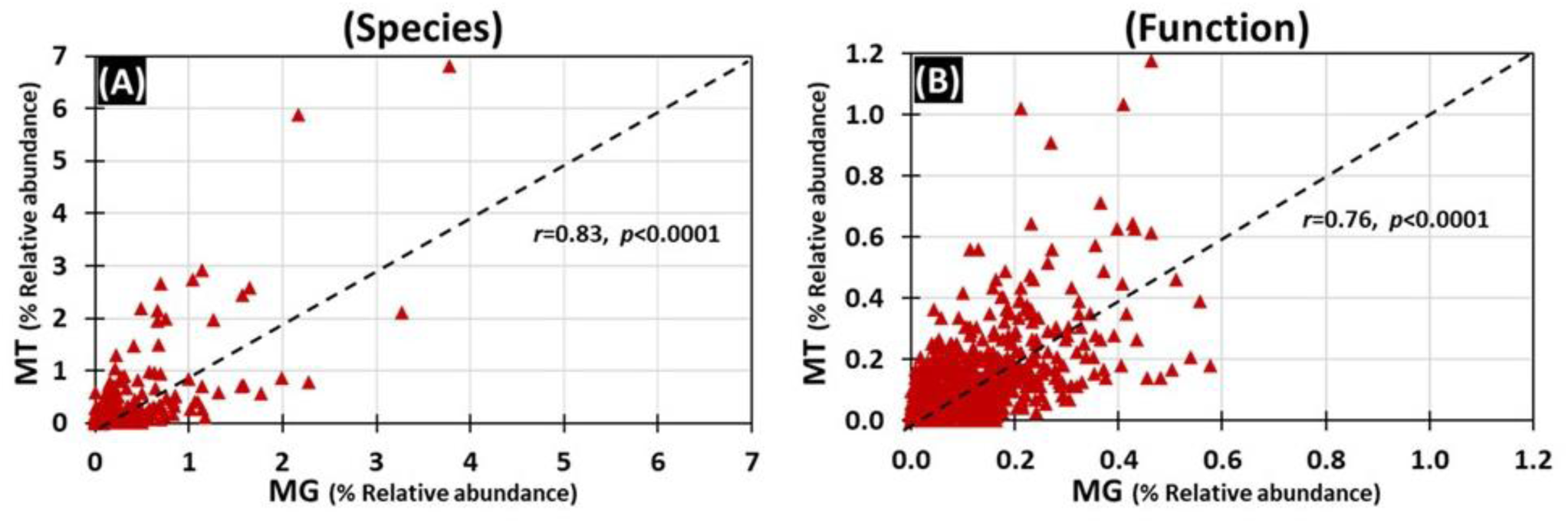
Correlation between the metagenome and metatranscriptome. Linear regression analysis was applied to the MT and MG data examined from the perspective of species and function. Spearman’s rank correlation between MG and MT (A) Bacterial species, (B) Functions at KEGG Level 4.

## DISCUSSION

Dysbiosis of the microbiome has been associated with multiple disease states including obesity, inflammatory bowel disease, asthma, urinary tract infection, cardiovascular disease and cancer (Pflughoeft & Versalovic 2012, Rani et al 2016a, Rani et al 2016b). However, the biological mechanisms that link the complex community of a microbiota with the pathogenesis of most diseases remains elusive. One limitation of many studies has been the use of targeted 16S rRNA amplicon sequencing which is generally limited to the genus and or OTU level of classification, thus, a more specific classification at the species level is not available (Metwally et al 2016, Metwally et al 2018). In contrast, MGS deep sequencing can accurately classify bacteria at the species level and also facilitates the annotation and identification of genes which predict putative biological functions. Further, due to the transcriptional regulation of many genes, MGS sequencing does not reveal gene expression levels. To address both the challenges, in this project we have optimized and evaluated the combination of metagenomic and metatranscriptomic shotgun sequencing data to evaluate methods to analyze the functional roles of both abundant and rare species in the microbiota. We generated 139.6 million metatranscriptomic reads which we compared to our previously reported metagenome shotgun sequencing data on the same sample that included 163.7 million reads (Ranjan et al 2016). One of the limitation of this study is sample size, as is it focused on n-of-1, and these findings may not be observed in different biological samples. However, with the advent of personalized medicine and clinical translational studies, there has been surge of n-of-1 studies. Many of clinical cases possess unique features that may not be identified by classical studies involving large number of samples (Nikles et al., 2010; Lillie et al., 2011; Schork, 2015).

First, our study shows that the different Illumina platforms do not contribute detectable bias in our analyses (Fig. 2). To validate the technical reproducibility of the sequencing and data analysis methods, we generated 12 replicates of a single sample that generated a similar number of reads, total bases and assembled contigs (Table 1). In addition, our analysis identified a reproducible number of both phyla and species (Supplementary Figs. 2 and 4, respectively). Furthermore, the functional analysis identified similar abundance of KEGG annotations at all functional levels from 1-4 (see Supplementary Figs. 6-9). Our investigation of the effect of contig assembly showed that assembly only modestly increased length,
presumably due to the short length of the mRNA transcripts. This similar observation has also been reported in a forest floor community metatranscriptomics (Hesse et al 2015). This suggests that emerging technologies that produce longer read lengths, particularly in view of their increased error rates, although useful for metagenomics studies, may not be preferable for metatranscriptomic studies.

Our investigation of the effects of contig assembly showed that the relative abundance of some taxa was modified by assembly. For example, analysis of assembled reads resulted in greater abundance of Bacteroidetes and lesser abundance of Firmicutes, Actinobacteria and Proteobacteria (Supplementary Fig. 5). Similar differences were also observed at the level of genus and species. Interestingly, we also observed similar changes in relative abundance of Bacteroidetes and Firmicutes in our previous analysis of taxa assignment in our metagenomics data (Ranjan et al 2016). Our results also show that the assembly of reads into contigs can decrease the detection of taxa. Overall, the results suggest that reads are the most comprehensive, and contigs are more specific, method to annotate taxa.

Most previous microbiota studies have not been performed with matched metagenome and metatranscriptome datasets of the same sample, thus there is huge knowledge gap in understanding the role of gene expression of the microbiota in human health and diseases. Our comparison of the predicted functions in the metagenome in this sample, with the expressed functions in metatranscriptome, identified more than 1000 functions, which included carbohydrate metabolism, nucleotide metabolism, amino acid metabolism, translation etc., (Fig. 5). The diversity analysis also suggest that the actual metabolically active bacterial species and functions are in fact less diverse compared to predicted metagenome diversity (both taxonomic and functional) (Supplementary Fig. 19).

It is well established that the diverse community of bacteria in a microbiome is composed of a small number of abundant species plus a large number of low or rare abundance species (Ranjan et al 2016); however, the functional role of the abundant versus rare species is not well understood. Our comparison of the metatranscriptome with the metagenome data suggests that both the abundant and rare bacteria may be actively engaged in the gut ecosystem. For instance, bacterial transcripts representing phyla Firmicutes (*F. prausnitzii*), and Bacteroidetes (*Bacteroides spp*., and *B. uniformis*) were highly abundant in MT (Fig. 3). Bacterial phyla - Fusobacteria and Verrucomicrobia are relatively less abundant in human gut, but are known to play an important role gut physiology (Everard et al 2013, Tremaroli & Backhed 2012). For instance, in our sample, both these phyla actively contributed in expression of specific genes involved in carbohydrate and amino acid metabolism pathways (Figs. 6 and 7). For example, genes such as *glgB* (1,4-alpha-glucan branching enzyme) and *pgi* (glucose-6-phosphate isomerase) involved in starch and sucrose metabolism and gluconeogenesis/glycolysis were highly expressed by Fusobacteria. These data suggest that the low abundant bacterial species are not just mere bystanders but actively contribute to the gut ecology. A similar study using the matched metagenomics and metatranscriptomics of the same sample have observed comparable findings that microbial and metabolic potential vary and are not concordant with their taxonomic abundance (Franzosa et al 2014). The functional potential of the more and less abundant bacterial species remain poorly understood. However, our observations indicate that the less abundant species are also metabolically active and may play unique roles in host-bacteria and bacteria-bacteria interactions and may actively contribute to the gut microbiota and physiology.

## ACKNOWLEDGEMENTS

This work was supported in part by NIH RO1 HL081663 and NIH RO1 AI053878 to DLP and PWF. The authors acknowledge Mr. Samer Sabbagh for help with preparing the libraries.

## AUTHOR CONTRIBUTIONS

DLP, PWF, RR and AR designed the study: RR prepared libraries and performed sequencing, AR and RR performed data analysis, RR, AR, PWF and DLP wrote the manuscript.

## SEQUENCE DATASETS

The sequence data files have been submitted to MG-RAST and the accession numbers are mentioned in Supplementary Table 5.

## SUPPLEMENTARY FIGURE AND TABLE LEGENDS

### Supplementary Figure 1. Fecal metatranscriptome library preparation

High quality total RNA from a fecal sample was isolated and analyzed by agarose gel electrophoresis and Bioanalyzer (A); Total RNA was enriched for mRNA by depleting the rRNA by subtractive hybridization method (B), the enriched mRNA was fragmented by Covaris (C); A library was prepared using Illumina compatible adaptor (D); In addition, 12 libraries from the same mRNA were prepared for multiplexing (E). The quality of RNA, mRNA and the libraries was analyzed on 2100 Bioanalyzer Instrument.

### Supplementary Figure 2. Phylum level analysis of multiplexed libraries using read and contig based analysis

The twelve metatranscriptome libraries were sequenced on Illumina MiSeq (151 PE) and analyzed for bacterial taxonomic assignment at phylum level using sequence read (A) and assembled contigs (B). Also, all the twelve libraries were combined in-silico and called as Lib-all.

### Supplementary Figure 3. Genus level analysis of multiplexed libraries using read and contig based analysis

The twelve metatranscriptome libraries were sequenced on Illumina MiSeq (151 PE) and analyzed for bacterial taxonomic assignment at genus level using sequence read (A) and assembled contigs (B). Also, all the twelve libraries were combined in-silico and called as Lib-all, and top 1% genus are shown (data sorted high to low abundance in Lib-all).

### Supplementary Figure 4. Species level analysis of multiplexed libraries using read and contig based analysis

The twelve metatranscriptome libraries were sequenced on Illumina MiSeq (151 PE) and analyzed for bacterial taxonomic assignment at species level using sequence read (A) and assembled contigs (B). Also, all the twelve libraries were combined in-silico and called as Lib-all, and top 1% bacterial species are shown (data sorted high to low abundance in Lib-all).

### Supplementary Figure 5. Taxonomic analysis of the metatranscriptome

The read and contig based analysis of HS100, MS151, MS301, and HS100-MS151-MS301 (A). (B) The MT for each sequencing strategy (HS100, MS151 and MS301) was sampled for 30M reads. The reads were assembled into contigs and analyzed for taxonomic annotations based on read and contig. Data is sorted high to low on MS301_read dataset.

### Supplementary Figure 6. Functional analysis of metatranscriptome at level 1 of multiplexed libraries using read and contig based analysis

The twelve metatranscriptome libraries were sequenced on Illumina MiSeq (151 PE) and analyzed for functional assignment at Level 1 using MGRAST KEGG module using sequence read (A) and assembled contigs (B). Also, all the twelve libraries were combined in-silico and called as Lib-all, and all the six Level 1 functions are shown.

### Supplementary Figure 7. Functional analysis of metatranscriptome at level 2 of multiplexed libraries using read and contig based analysis

The twelve metatranscriptome libraries were sequenced on Illumina MiSeq (151 PE) and analyzed for functional assignment at Level 2 using
MGRAST KEGG module using sequence read (A) and assembled contigs (B). Also, all the twelve libraries were combined in-silico and called as Lib-all, and top 10 Level 2 functions are shown. The data is sorted high to low on Lib1.

### Supplementary Figure 8. Functional analysis of metatranscriptome at level 3 of multiplexed libraries using read and contig based analysis

The twelve metatranscriptome libraries were sequenced on Illumina MiSeq (151 PE) and analyzed for functional assignment at Level 3 using MGRAST KEGG module using sequence read (A) and assembled contigs (B). Also, all the twelve libraries were combined in-silico and called as Lib1 −12, and top 10 Level 3 functions are shown. The data is sorted high to low on Lib1.

### Supplementary Figure 9. Functional analysis of metatranscriptome at functional level 4 of multiplexed libraries using read and contig based analysis

The twelve metatranscriptome libraries were sequenced on Illumina MiSeq (151 PE) and analyzed for functional assignment at Level 4 using MGRAST KEGG module using sequence read (A) and assembled contigs (B). Also, all the twelve libraries were combined in-silico and called as Lib1 −12, and top 1% Level 4 functions are shown. The data is sorted high to low on Lib1.

### Supplementary Figure 10. Functional analysis of metatranscriptome based on read and contig

(A) Level 1, (B) Level 2, (C) Level 3, and (4) Functional. The MT for each sequencing strategy (HS100, MS151 and MS301) was sampled for 30M reads. The reads were assembled into contigs and analyzed for taxonomic annotations based on read and contig. Data is sorted high to low on MS301_read dataset. For Level 1 all functional categories are shown, for Levels 2-4, only top 10 functions are shown.

### Supplementary Figure 11. Abundance of bacterial species in different phyla in MG and MT

Abundance of bacterial species in different phyla - Firmicutes, Bacteroidetes, Actinobacteria, Proteobacteria, Fusobacteria and Verrucomicrobia. Note the higher abundance percentage in metatranscriptome compared to metagenome data, indicating that some species are more metabolically active. Only top 10 species are shown for MT-HS100-MS151-MS301 and MG-HS100-MS151-MS301 (data sorted on MT-HS100-MS151-MS301).

### Supplementary Figure 12. Functional analysis at level 1

Percent abundance of the predicted (based on metagenome) and expressed (metatranscriptome) function. Data is sorted high to low on MT-HS100+MS151+MS301.

### Supplementary Figure 13. Functional analysis at Level 2

Percent abundance of the predicted (based on metagenome) and expressed (metatranscriptome) function. Functions are sorted high to low on MT-HS100+MS151+MS301 and above 1% are reported.

### Supplementary Figure 14. Functional analysis at Level 3

Percent abundance of the predicted (based on metagenome) and expressed (metatranscriptome) function. Data is sorted high to low on MT_HS100+MS151+MS301, and top 10 functions and above 1% are reported.

### Supplementary Figure 15. Functional analysis at functional level

Percent abundance of the predicted (based on metagenome) and expressed (metatranscriptome) function. Functions is sorted high to low on MT_HS100+MS151+MS301 and top 10 functions are reported.

### Supplementary Figure 16. Functional analysis at level 1 in individual phylum

The functions in individual phylum were analyzed in the metatranscriptome (MT-HS100+MS151+MS301) data.

### Supplementary Figure 17. Functional analysis at level 2 in individual phylum

The functions in individual phylum were analyzed in the metatranscriptome (MT-HS100+MS151+MS301) data.

### Supplementary Figure 18. Functional analysis at level 3 in individual phylum

The functions in individual phylum were analyzed in the metatranscriptome (MT-HS100+MS151+MS301) data and sorted high to low on Firmicutes and above 1% functions are shown.

### Supplementary Figure 19. Diversity indices for bacterial species (A) and functions (B), in MG and MT

The Shannon diversity and evenness are calculated for MG using the contig assembly of data MG-HS100, MG-MS151, MG-MS301 and MG-HS100+MS151+MS301, and MT using the contig assembly of data MT-HS100, MT-MS151, MT-MS301 and MT-HS100+MS151+MS301.

### Supplementary Figure 20. KEGG metabolic pathway in metagenome and metatranscriptome

Functions identified in the metagenome (MG-HS100+MS151+MS301) and metatranscriptome (MT-HS100+MS151+MS301). Blue: predicted functions exclusive in metagenome; Red: Exclusive in metatranscriptome.

### Supplementary Table 1

List of bacterial species identified based on read based analysis. Only above 1% are mentioned and sorted high to low on Lib1.

### Supplementary Table 2

List of bacterial species identified based on contig based analysis. Only above 1% are mentioned and sorted high to low on Lib1.

### Supplementary Table 3

Random sampling of the metatranscriptome sequence read and de-novo assembly of contigs.

### Supplementary Table 4

Abundance of bacterial species in metatranscriptome data based on read and contig analysis.

### Supplementary Table 5

List of accession numbers.

### Supplementary Table 6

List of bacterial species/sequences identified in the metatranscriptomics data.

